# Comparison of germline and somatic structural variants in cancers reveal systematic differences in variant generating and selection processes

**DOI:** 10.1101/2023.10.09.561462

**Authors:** Wolu Chukwu, Siyun Lee, Alexander Crane, Shu Zhang, Sophie Webster, Ipsa Mittra, Marcin Imielinski, Rameen Beroukhim, Frank Dubois, Simona Dalin

## Abstract

Although several recent studies have characterized structural variants (SVs) in germline and cancer genomes, the features of SVs in these different contexts have not been directly compared. We examined similarities and differences between 2 million germline and 115 thousand tumor SVs from a cohort of 963 patients from The Cancer Genome Atlas (TCGA). We found significant differences in features related to their genomic sequences and localization that suggest differences between SV-generating processes and selective pressures. For example, we found that transposon-mediated processes shape germline much more than somatic SVs, while somatic SVs more frequently show features characteristic of chromoanagenesis. These differences were extensive enough to enable us to develop a classifier-“the great GaTSV”-that accurately distinguishes between germline and cancer SVs in tumor samples that lack a matched normal sample.

## Introduction

Structural variants (SVs) are rearrangements of genomic material that result from incorrect double-strand break (DSB) repair. Large-scale whole-genome sequencing (WGS) efforts have revealed a prominent role of recurrent SVs as cancer drivers^1^ and as biomarkers of disruption of the DNA damage response and other processes^2,3^. Similar studies on normal tissue have shown that human genetic diversity results in large part from germline SVs^4^. The SVs in these different contexts may result from different biological constraints, yet germline and cancer SVs have not been directly compared.

SVs arise from DSBs that are repaired to incorrect loci in the genome. The resulting rearrangements can delete, duplicate, invert, or translocate segments of DNA. The responsible mechanisms of DSB repair include non-homologous end joining (NHEJ), which pastes broken ends of DNA together irrespective of adjacent sequence homology; microhomology-mediated end joining (MMEJ), which utilizes 3-10bp of breakpoint-adjacent microhomology to form repair intermediates; and homologous recombination (HR), which uses more bases of homology.

Germline and somatic SVs are likely to arise from different DSB repair mechanisms. Previous studies report that germline SVs primarily result from non-allelic homologous recombination (NAHR), where substantial amounts of sequence homology at distinct loci are employed in double-stranded break repair, often resulting in the deletion of the intervening sequence ^5,6^. Conversely, somatic SVs, which present with more varied spans and clustering of breakpoints, indicate a tendency towards NHEJ and replication-based mechanisms of repair such as microhomology-mediated break-induced replication (MMBIR) which are more error-prone^5–7^. However, a direct and comprehensive comparison between the features of germline and somatic SVs is likely to provide a more detailed view of the differences in their generation, with implications for the activity of different DNA damage and repair processes in human vs. somatic cell evolution.

One benefit of recognizing the different features of germline and somatic SVs is that it provides an opportunity to distinguish germline and somatic SVs when only data from somatic tissue have been collected. Currently, confidently designating an SV as “somatic” requires comparing sequencing data from highly clonal somatic tissue such as cancers or single cells with multiclonal “normal” tissues that represent the germline. However, there are many situations in which normal tissue is unavailable, including clinical settings^8^ and the study of long-term cell line models^9^. The ability to distinguish germline from somatic SVs in the absence of normal tissue is therefore valuable both clinically and in cancer research.

Here, we comprehensively evaluate similarities and differences between germline and somatic SVs. We find that they are strikingly different, enabling us to develop a machine learning-based classifier (the <u>G</u>ermline <u>a</u>nd <u>T</u>umor <u>S</u>tructural <u>V</u>ariant classifier, aka great GaTSV) that can distinguish germline from somatic SVs in the absence of a matched normal sample with extremely high accuracy.

## Results

To explore the differences between somatic and germline SVs, we used a TCGA dataset of paired tumor-normal WGS encompassing 963 tumors from 24 cancer types. We used the SvABA SV caller to ascertain the breakpoints and types of the SVs in this dataset^10^. Across the 963 tumors, germline SVs outnumbered somatic SVs 17:1 (**Fig. 1A**, median of 2,007 germline SVs and 53 somatic SVs per tumor). The number of germline events in a given individual was constant irrespective of age, whereas the number of somatic SVs in a sample showed a slight positive correlation with age (**Fig. 1B-C**).

**Figure 1.**
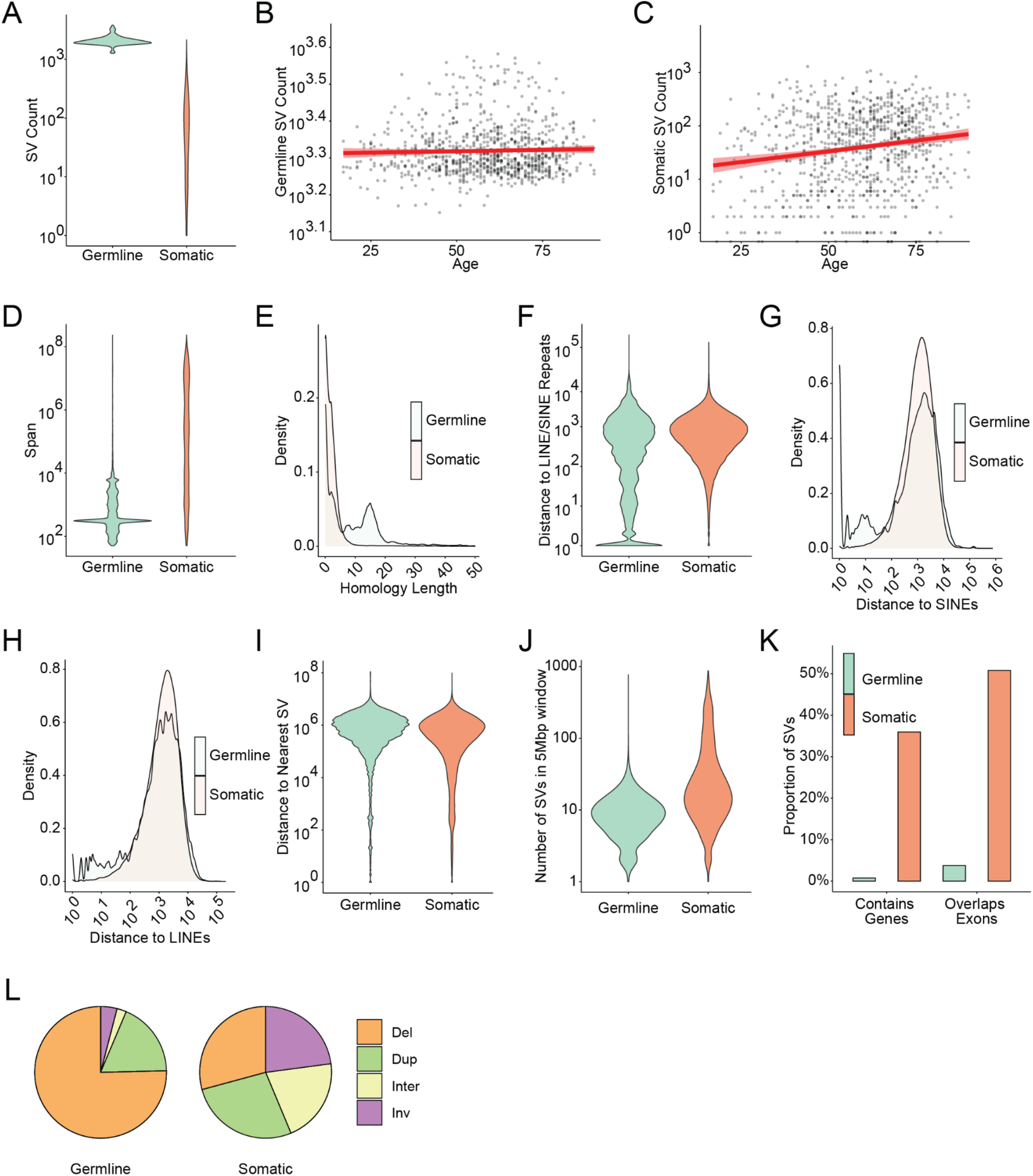
SV features differ significantly between germline and somatic genomes. **A**, Distribution of germline and somatic SV frequencies in TCGA genomes. **B-C**, Pearson correlation analysis between germline SV frequency and patients’ age in germline (B) and somatic (C) genomes. **D**, Overall span distribution of germline and somatic SVs. **E,** Homology length distribution of germline and somatic SVs. **F**, Distance of germline and somatic SVs to nearest repeat element (LINE or SINE). **G-H**, Distribution of distance to SINE (G) and LINE (H) elements. **I**, Distance to nearest SV within a sample. **J**, Number of SVs within a 5Mbp window of each SV breakpoint within a sample. **K**, Proportion of germline and somatic SVs that impact a gene or overlap an exon. **L**, Proportion of each SV type-Del(deletion), Dup(duplication), Inter(interchromosomal/translocation), Inv (inversion) present in germline and somatic SVs.

### Germline and somatic SVs have different SV features

To evaluate the differing impacts of SV generation and selection processes between germline and somatic contexts, we compared features of germline vs. somatic SVs (also see **Supplementary Table 1 and Supp. Fig S1A-D**). The most striking difference was SV span, the distance between intrachromosomal breakpoints. Somatic SVs had spans 60 times larger than those of germline SVs (KS test, p-value < 2.2e-16; **Fig 1D**) and were about twice as likely to have spans greater than 1,000 bp (67% of all somatic SVs vs. 31% of all germline SVs), a number that swells to 60 times more likely at 1 Mb (27% of all somatic SVs vs. 0.4% of all germline SVs). These results align with the intuition that SVs of larger spans are likely to result in changes to the genome that are not tolerated during normal development. Peaks in the germline SV span distribution corresponded to the typical spans of SINE and LINE transposable elements^11,12^. Germline and somatic span distributions varied by SV type, with the greatest differences between germline deletions, which tend to be short, and somatic deletions, whose span distribution is more uniform (**Supp Fig. S1E**). These results are consistent with previous studies of germline and somatic SVs ^2^.

The second most striking difference between germline and somatic SVs was the much higher levels of breakpoint homology attributed to germline SVs (KS test, p-value < 2.2e-16; **Fig 1E**), suggesting a transposon-mediated origin. Closer examination of the distribution of homology lengths revealed a peak between 13-17bp in germline SVs but not somatic SVs. This peak was specifically present in germline deletions of ∼300bp span (**Supp Fig. S1F-G**), corresponding to the spans of Alu elements-a type of SINE element and the most abundant transposons in the human genome^13^. Previous studies have shown that Alu elements comprise a significant proportion of transposable element-mediated rearrangements in the genome and use ∼15 bp of homology^14^. Germline SVs were closer to SINE and LINE elements than somatic SVs (KS-test, p-value < 2.2e-16, **Fig. 1F-H**, **Supp Fig. S2A-B**), and—of all repeat elements—these showed the greatest difference in range of distances between germline SVs and somatic SVs (**Supp Fig. S2C-Q**). Interestingly, somatic SVs were closer to all classes of RNA pseudogenes compared to germline SVs.

In contrast, somatic SVs are more likely to be formed by chromoanagenesis^15^ and affect gene structure. Somatic SVs were more likely to be found in proximity to each other (**Fig. 1I-J**, p-value < 2.2e-16, KS-test) and were more likely to disrupt coding sequences or span entire genes (Fisher’s Exact Test, both p-values < 2.2e-16, **Fig 1K**). Strikingly, 51% of somatic SVs directly affected the exome, in contrast to only 3.8% of germline SVs. Finally, deletion events made up about 75% of germline events but only 29% of somatic events (**Fig 1L**, Chi-Squared test, p-value < 2.2e-16), whereas somatic SVs were nine times more likely to be translocations (Chi-Squared test, p-value < 2.2e-16). Overall, our analysis of single features of SVs showed pronounced differences in the characteristics of germline compared to somatic SVs.

If there are specific variant-generating or selection processes that result in germline and somatic SVs, we expect the features of SVs (**Supplementary Table 1**) that are linked to such processes to vary together. Indeed, characteristic combinations of SV features between germline and somatic SVs reflected the pronounced differences in the relationships between SV features (**Fig. 1A-C**). Somatic SVs with shorter spans tended to have longer homology (**Fig. 1A**), while no such associations were observed for germline SVs. Germline SVs had longer homology if they were closer to known SINE elements or were deletions (**Fig. 1A, 2B**), while no such association existed for somatic SVs (**Fig. 1A, 2D**). Longer germline SVs also showed higher homology GC content while we observed the opposite in somatic SVs (**Fig. 1D**). These data suggest that transposon-mediated processes dominate germline SV generation while only a subset of somatic SVs originate from this pathway. In contrast, somatic SVs showed an anti-correlation between homology length and GC content with the number of SVs within 5Mbp as well as SV span with its distance to the nearest SV (**Fig 2A, 2D**). These data indicate a subset of (longer) somatic SVs generated in clusters by NHEJ in a more complex process like chromothripsis.

**Figure 2.**
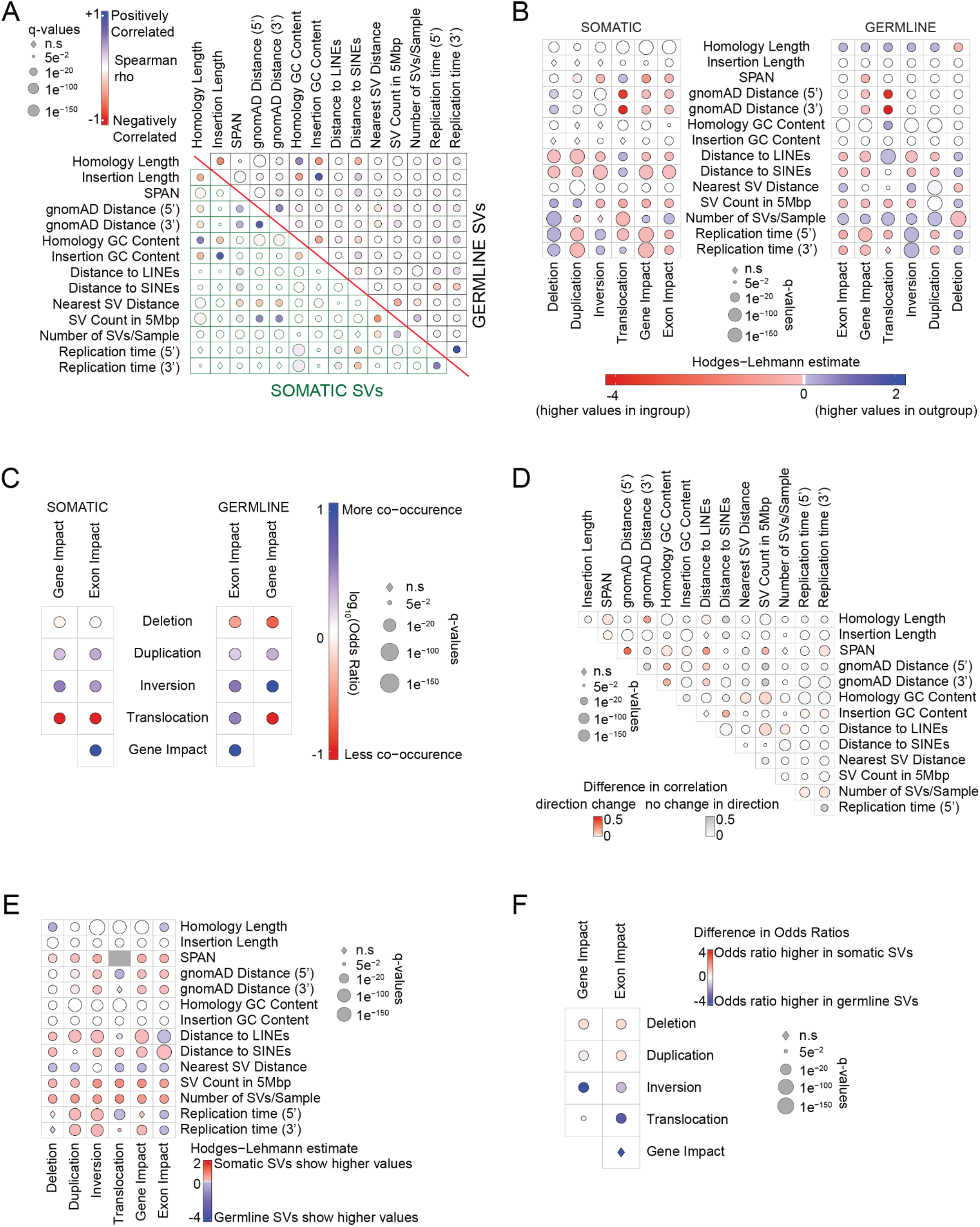
SV feature associations differ between somatic and germline SVs. **A**, Spearman correlations between continuous features within somatic and germline SVs (positive correlations in blue, anticorrelation in red). **B**, Differences in the values of continuous SV features between categorical SV features within germline and somatic SVs. **C**, Odds ratios of the associations between the categorical variables gene or exon impact and SV type in germline and somatic SVs. **D**, Difference in Spearman correlation values between continuous features across germline and somatic SVs. **E**, Differences in the values of continuous SV features between germline and somatic SVs within categorical SV features. **F**, Difference in odds ratios of categorical variable comparisons across germline and somatic SVs. For all panels, the circle size represents the significance of the statistic represented.

The differences in feature associations between germline and somatic SVs also hinted at the strong differences in the selection pressures they face. Somatic deletions were significantly more likely to span gene/exons than germline deletions. (**Fig 2C, F**). Also, while duplications were also more likely to span genes than other SV types in both germline and somatic SVs (**Fig. 1C, q**-value < 2.2e-16, Chi-Squared test), this association was significantly stronger in somatic SVs (**Fig. 1F**). Germline inversions, which could be more neutral in their effects on gene expression, were more likely to affect genes/exons than other germline SV types (**Fig. 1C**) and also more likely to span genes/exons than somatic inversions (**Fig. 1F)**. Our analysis of the association between replication timing and SV type also found results consistent with strong selection pressures conserving the coding genome in the germline. We observed a positive correlation between homology length and replication timing among germline SVs that was absent or insignificant among somatic SVs (**Fig. 1A**). This shows an enrichment for high-fidelity repair in gene-dense regions in the germline context that is not observed in cancer-associated SVs. Also, while there was an overall bias for inversions and deletions towards later replicating genomic loci and duplications for earlier replicating loci (**Fig. 1B**), striking differences arise when comparing across germline and somatic SVs. Notably, somatic duplications and inversions showed significantly higher replication timing values, corresponding to earlier replicating regions, than germline duplications and inversions (**Fig. 1E**). Early-replicating regions tend to be gene-dense and highly expressed.^16^ Together, these data indicate strong selection pressures to conserve the full coding genome in the germline that are attenuated in cancer cells.

### Extending SNV classification approaches to SVs

Next, we considered how we might use this information to distinguish between germline and somatic SVs when paired germline DNA sequencing data are unavailable. As a control, we first attempted to filter germline SVs using the gnomAD v4.0 population dataset^17^, analogous to the commonly-used filtering approach for removing germline SNVs^18,19^. However, only a fraction of germline SVs matched within 3bp of a gnomAD SV (∼1.0M SVs out of 2.0M germline SVs with only ∼31k SVs matching exactly to a reference gnomAD SV). The paucity of exact matches may be due to differences in SV callers between gnomAD and our dataset as well as the fact that the vast majority of germline SVs are not recurrent^17^. While the distance from gnomAD-listed germline SVs was significantly different for germline and somatic SVs (**Supp. Fig S3A**, p-value < 2.2e-16, **Supp. Fig S3A**), about a third of germline SVs lay more than 1000bp from a gnomAD SV. Therefore, to match TCGA SVs to the gnomAD reference SVs, we determined the average base pair distance of each pair of breakpoints from each TCGA SV to its closest SV in gnomAD. This “fuzzy matching” approach had an overall AUC of 0.90. The optimal cutoff-point on the ROC curve of ∼1400bp average distance from the nearest gnomAD SV resulted in a high degree of sensitivity in correctly classifying somatic SVs (TPR ⋍ 96% of true somatic events). However, this cutoff still resulted in a large fraction of germline contamination (60%) in the set of SVs called somatic (positive predictive value, PPV) (**Fig 3A)**. These data show that a simple filter based on the closest germline SV in gnomAD cannot sufficiently differentiate between germline and somatic SVs in tumor-only SV calls.

**Figure 3.**
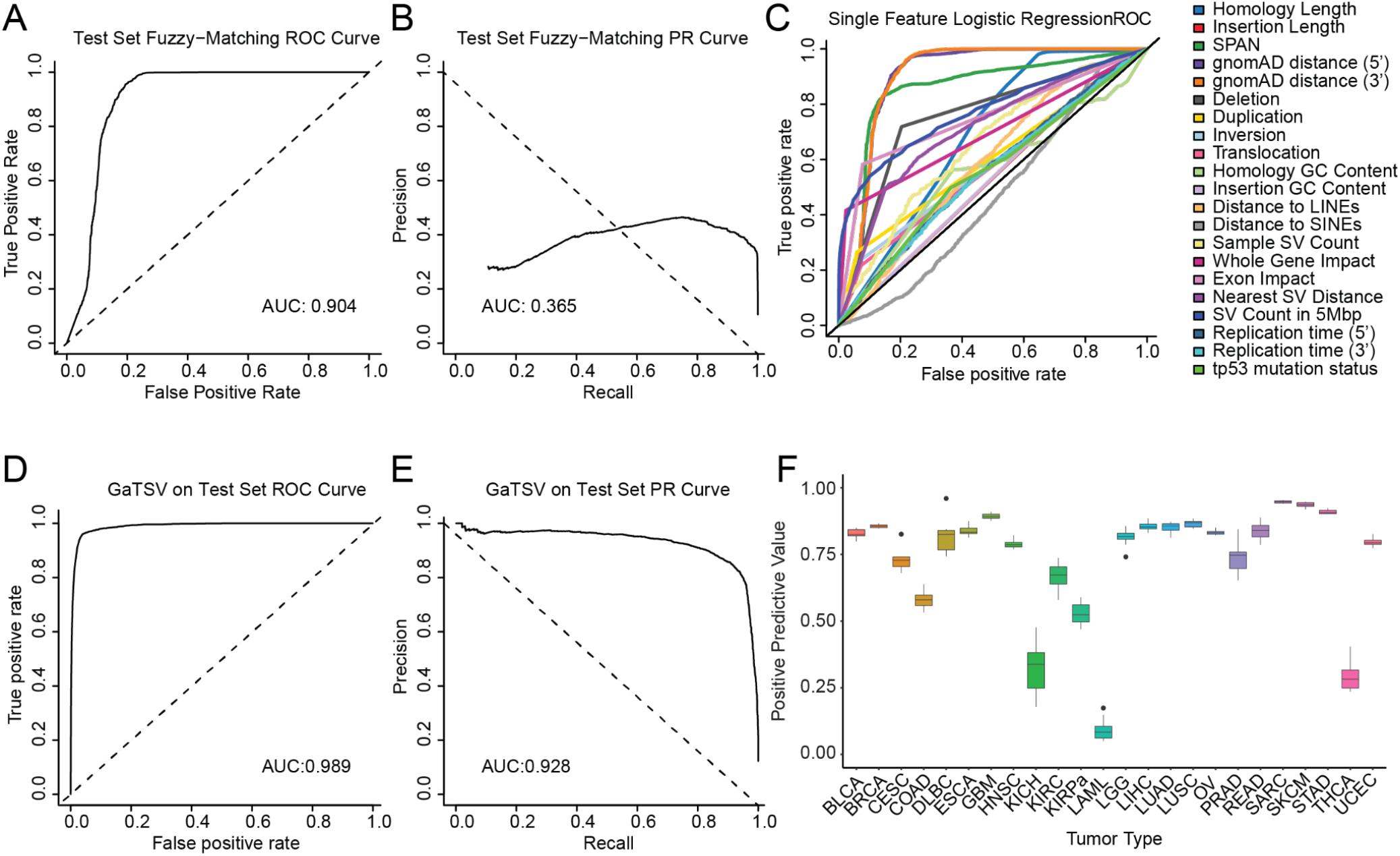
Classifier performances on TCGA test set. **A**, ROC for classifying SVs based on a cutoff distance to gnomAD reference (fuzzy-matching). **B**, PR curve for classifying SVs using fuzzy-matching. **C**, ROC for separate single-feature logistic regression classifiers. **D**, ROC for the GaTSV classifier. **E**, PR curve for the GaTSV classifier. **F**, Positive Predictive Value (PPV) of the GaTSV classifier by tumor type showing variance in performance across tissue types.

**Figure 4.**
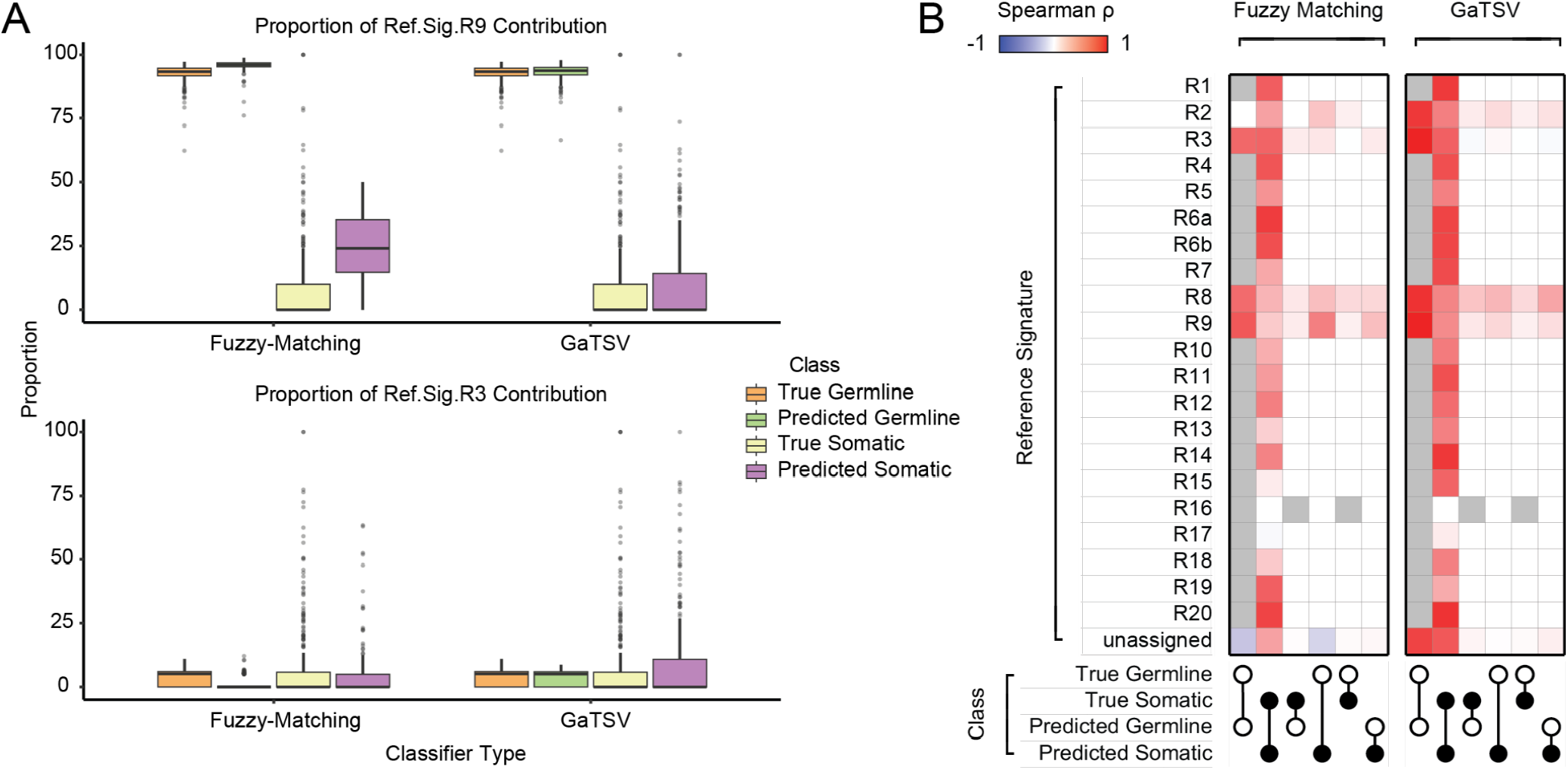
Analysis of classifier performances using rearrangement signatures. **A**, R3 and R9 reference signature activity is more accurately recapitulated using GaTSV classified SVs than fuzzy-matching classified SVs. The proportion of contributions of the R9 reference signature (top) and R3 reference signature (bottom) in each patient. Box colors represent whether the germline or somatic SVs calls originated from SvABA (true), GaTSV, or fuzzy-matching (predicted). **B**, Signature contributions based on GaTSV classified SVs more closely match the true distributions compared to the fuzzy-matching classified SVs. Similarity of signature contributions based on true SVs and SVs called by each classifier were calculated by Spearman correlation for all reference signatures between all combinations of predicted vs. true germline vs. somatic SVs. The gray squares refer to undefined correlations between two zero vectors, or when no samples in either class have any contribution for a given reference signature.

### Single features are insufficient to distinguish germline and somatic SVs

To test the predictive value of individual SV features, we constructed single-feature logistic regression models with a training set of 200,000 SVs. We tested each model on 100,000 SVs and obtained an average single-feature AUC of 0.656 (SD 0.127, **Fig 3C)**. The distance from either breakpoint of an SV to its nearest gnomAD SV performed best, with similar AUCs of ∼0.90 (**Supplementary Table 2**). Even with this performance, neither feature achieved a positive predictive value of greater than 0.50 resulting in a large amount of germline SVs in the set of predicted-somatic events. Replication timing, insertion GC%, insertion length, and distance to SINE elements were poor individual differentiators between germline and somatic SVs. We concluded no single SV characteristic was sufficiently predictive to perform the germline or somatic classification.

### Classifier Development and Performance

As individual features cannot sufficiently distinguish germline and somatic SVs, we used the combination of SV features to develop a support vector machine (SVM) based classifier-the great <u>G</u>ermline <u>a</u>nd <u>T</u>umor <u>S</u>tructural <u>V</u>ariant (GaTSV) classifier-to distinguish between germline and somatic SVs. We trained GaTSV on 509,433 SVs from 634 samples, two-thirds of all samples available in our TCGA cohort. Once trained, we tested the classifier on 262,118 SVs from 329 samples, one-third of the samples in our TCGA cohort.

In addition to a binary classification, GaTSV generates a probability of classification for each variant, with values closer to 1 representing a higher likelihood of a variant being somatic. These probabilities allow for the selection of a classification cutoff to prioritize certain performance metrics, including AUC, PPV, etc. For instance, having a cutoff farther from 0 would reduce the number of germline SVs falsely called somatic by the classifier (increasing the PPV), while increasing the number of true somatic SVs falsely called germline (decreasing the TPR). To balance these considerations, GaTSV uses a cutoff of 0.268 which maximizes the sum of the PPV and TPR in our test set.

Overall, GaTSV achieved an AUC of 0.989, with a sensitivity (TPR) of 0.915, specificity (1-FPR) of 0.977 (**Fig. 1D-E**), and PPV of 0.849.

GaTSV’s performance varied across tumor types, largely due to differences in the prevalence of somatic SVs in each tumor type. The fraction of GaTSV-classified somatic SVs that were truly somatic was higher in tumors with many somatic SVs. In other words, the PPV was higher in tumors with more somatic SVs. In our test set, sarcomas had the highest somatic SV burdens (∼391 SVs per tumor) and the highest proportion of called-somatic SVs that were truly somatic (PPV = 0.947). In contrast, acute myeloid leukemia, with the fewest SVs per tumor (∼2 SVs), had the lowest proportion of called-somatic SVs that were truly somatic (0.09) (**Fig. 1**). This finding was corroborated at the sample level. Samples with a lower SV burden in the test set tended to have lower PPV than samples with higher SV burden (**Supp Fig. S3B-C**). Specifically, 86% of samples with fewer than 10 somatic SVs had PPV below 0.5. In contrast, only 0.5% of samples with 10 or more SVs had a PPV below 0.5 (p-value < 2.2e-16, Fisher’s Exact Test). These data show that GaTSV performs well overall but is subject to falsely calling true germline SVs as somatic SVs in samples with very low somatic SV counts.

GaTSV achieved an accuracy of 97% on average across all tumor types (**Supp Fig. S3D**). To uncover potential weaknesses of our classifier, we examined the features of the misclassified SVs. The feature distribution of germline SVs mislabeled as somatic largely mimics that of true somatic SVs in the test set. The patterns of correlations between the SV features also resemble true somatic SVs more than true germline SVs (**Supp. Fig. S4**). This suggests that GaTSV poorly differentiated these somatic-like germline events, leading to germline contamination in the somatic calls. For example, 98% of misclassified germline SVs were more than 1kb away from a gnomAD reference (**Supp. Fig. S4B**). Nonetheless, GaTSV correctly classified 91% of germline SVs with a distance of 1kb or more from a gnomAD SV; therefore, GaTSV performs well on most SVs.

### GaTSV’s performance in an Independent Dataset

To test how well GaTSV classifies SVs in tumors from other datasets, we gathered a test set consisting of 7,623 SVs from six pediatric high-grade glioma (pHGG) patients with comparable somatic SV burden to TCGA samples. The tumor samples used in this dataset were collected with blood normals, which provided a truth set of somatic and germline SVs. The features observed in the pHGG dataset had similar distributions to those of the tumors in the TCGA dataset (**Supp**. **Fig. S5**). GaTSV achieved a sensitivity (TPR) of 0.975 and specificity (1-FPR) of 0.892. Of the SVs that were called somatic in this set, 84% were truly somatic (PPV: 0.839). We conclude that GaTSV performs robustly across different datasets.

### GaTSV’s performance across Different Ancestries

We hypothesized that imbalance in representation of individuals from certain ancestries in our training set would impair GaTSV’s performance on individuals of those ancestries-specifically, over 77% of individuals in the TCGA dataset are of European descent. Indeed, GaTSV performed better on SVs from European individuals than on SVs from East Asian and African individuals on all metrics considered, including the AUC of the ROC and PR curves as well as the PPV (**Supp Fig. S6A-F**). Among the ancestries considered, GaTSV performed the worst on African SVs across all metrics. African genomes are considerably more diverse than any other ancestry group, which means that increased representation in the training set is necessary compared to other ancestries.

To evaluate whether this drop-off in performance was unique to GaTSV, we also tested the gnomAD fuzzy matching method on different ancestries. Again, we saw the best performance on European SVs and the worst on African SVs, across all metrics (**Supp Fig. S6G-L**). European genomes are the most represented ancestry group in the gnomAD database at over 77%^20^. These results highlight the importance of ancestry biases in public databases. GaTSV’s performance on the worst-performing ancestry group—African—with a PPV of 0.66 was still notably better than the gnomAD fuzzy matching performance with a PPV of 0.42 on the best-performing ancestry group—European.

### GaTSV uncovers associations between somatic SV signatures and genetic variants in the absence of paired normal SV calls

The effects of somatic SV-generating processes were recently characterized across TCGA using SV signature analysis^3^. Somatic SV signature analysis has only been described after the removal of all germline SVs from the call set through joint analysis with paired normal tissue. We wanted to test if GaTSV allowed the accurate extraction of these biologically relevant patterns of somatic SVs without a paired normal. Organizing the set of all SVs in the TCGA dataset into the SV signature input catalog showed an abundance of short non-clustered deletions (**Supp Fig. S7A**), the majority of which were germline SVs. Both the GaTSV-classified germline SVs and the gnomAD fuzzy matching-classified germline SVs recaptured this true germline SV deletion peak. However, gnomAD fuzzy matching misclassified many of these short non-clustered deletions as somatic, resulting in a falsely inflated total number of somatic SVs to more than double the true count of somatic SVs. In contrast, GaTSV correctly matched both the distribution and counts of SVs seen in the true somatic and germline SV signature input catalogs.

We extracted the published SV signatures^3^ based on these catalogs and primarily detected reference SV signature R3 in the germline profile and only minor contributions of the other signatures. In contrast, somatic SVs were composed of a variety of signatures (**Fig. 1A**, **Supp Fig. S8A-B**). This difference in the reference signature proportions across germline and somatic SVs suggests that the mechanisms behind somatic SV generation are more varied than those behind germline SV generation. It is also possible that there are additional germline SV signatures that we did not detect because the reference signatures we used were extracted from somatic SVs.

In the comparison of reference SV signature proportions, the GaTSV also outperformed gnomAD fuzzy matching. The medians of gnomAD fuzzy matching derived “somatic” signature contributions were closer to the true germline than the true somatic contributions for reference signatures R3 (nonclustered translocations) and R9 (short inversions). In contrast, the GaTSV-predicted “somatic” signature contribution medians matched the respective true class medians (**Fig. 1A**). GaTSV also showed high levels of correlation for called somatic to true somatic and called germline to true germline for almost all reference signatures (**Fig. 1B**). These results further support our finding that the GaTSV overcomes the germline contamination issue in somatic SV calls without a paired normal.

## Discussion

Our analyses show that germline SVs are shorter, less likely to impact genes, and have more bases of homology adjacent to their breakpoints than somatic SVs. In contrast, somatic SVs are more likely to cluster together and are farther from transposable elements than germline SVs. These results strengthen previous findings that homology and transposon-based repair contribute more to the formation of germline SVs than somatic SVs. They also strengthen the intuition that germline genome structure is under much more stringent fitness constraints than somatic genome structure. We also establish that these differences can be used to computationally classify germline from somatic SVs in the absence of a matched normal, with high sensitivity and specificity.

Differences between DSB repair processes in somatic cells vs. germ cells and their progenitors are reflected in differences between SVs in these different contexts. Germline SVs tend to have more bases of homology than somatic SVs, revealing a preference for more accurate repair processes to maintain genome integrity. Somatic SVs are closer to each other than germline SVs, indicative of a higher likelihood of chromoanagenesis events in cancer cells.

Differences in fitness constraints in germline vs. somatic contexts are also reflected in their SVs. Most common germline SVs are short unclustered deletions without impact on a coding sequence. Previous work showed that even small repeats resulting from duplication have been found to decrease cell fitness^21^. Therefore, most germline SVs lack gene dosage effects or changes in protein composition. Ostensibly, such a rearrangement typically does not impact transcriptional regulation. Similarly, most germline SVs tend to accumulate close to previously described germline loci^17^ and are less likely to form in clusters. We postulate that these observations are the result of a selective process that eliminates de novo germline rearrangements that impact gene dose or protein functionality. The particular underlying process is worthy of further investigation and is plausibly attributed to embryonic lethality leading to variant extinction. As cells harboring somatic SVs do not need to develop from a single cell to an entire organism, they are likely subject to fewer fitness constraints than germline SVs. As a result, they are more tolerable to impacts on coding sequences and transcriptional dysregulation. We surmise that these SVs are under selection pressure that is divergent from germline SVs.

Previous studies have shown that the tumor context is capable of altering the replication timing landscape. Such transformation from early-to-late replication and vice versa are evidenced in chromatin remodeling and methylation frequency^16^. The types of SVs were also linked to this altered chromatin compaction density between germline and somatic cells. While this consideration is not accounted for in our analysis, future studies can seek to uncover this potential bidirectional relationship between the cancer context and replication timing.

We took advantage of the differences between germline and somatic SVs to create an SVM classifier to distinguish them in cases where a matched normal sample is unavailable. Our SVM performed well (AUC = 0.99, PPV = 0.85), so we termed it the great <u>G</u>ermline <u>a</u>nd <u>T</u>umor <u>S</u>tructural <u>V</u>ariant classifier-the great GaTSV classifier. One significant strength of GaTSV is its ability to classify SVs that are not present in germline reference databases-this is critical as most SVs are rare and therefore not present in reference databases. Our tool has several limitations. First, it did not perform as well on patients of African ancestry compared to patients of European ancestry, likely due to the underrepresentation of individuals of African descent in the TCGA training dataset we used. Second, our tool can only be used with SV calls from SvABA^10^ because it was trained on breakpoints called by SvABA, which may follow different conventions from other SV callers.

Another potential limitation of our study is the presence of artifactual rearrangements in SV calls. SvABA uses short-read sequencing data, potentially resulting in reads that each align to multiple repetitive loci. These multi-mapping reads can result in artifactual SV calls present in many germline and somatic samples. Since these SVs are present in both germline and somatic samples, they will be called germline by traditional methods that compare somatic SVs with a matched germline sample, such as our training set. Therefore, the GaTSV classifier will classify these artifactual SVs as germline. Hence, while we expect that the GaTSV classifier can identify somatic variants from the pool of germline and artifactual SVs, it is more difficult to draw biological conclusions from the GaTSV germline calls, as they are contaminated with artifactual SV calls. In future work, these issues can be addressed with long-read sequencing data, diversifying the ancestries represented in the training set, and extending GaTSV to use SV calls from other software such as MANTA and GRIDSS2^22,23^.

Although our classifier performed well overall, it tended to misclassify certain variants, such as its propensity to call somatic deletions shorter than 100kb as germline rearrangements. These incorrect classifications may reflect true limitations in the great GaTSV and create a false identity for SVs. However, our analysis showed that features of germline SVs misidentified as somatic closely mimicked features of true somatic SVs and vice versa. While SV-generating processes differ substantially between germline and somatic contexts, some processes such as the integration of transposable elements and repeats into the genome are common to both^24^. This overlap could lead to similarities in the SV features, thus making them indistinguishable to the classifier.

The great GaTSV classifier will facilitate several avenues of new research to understand the formation and impact of germline and somatic SVs. Using GaTSV, SVs in clinical samples lacking a matched normal can be interrogated. The power of deeply characterized groups of cell lines such as those in the Cancer Cell Line Encyclopedia^9,25,26^ can now be brought to bear on SVs, including discovering relationships between drug sensitivities, CRISPR dependencies, and gene expression on specific SVs, signatures of SVs, and SV abundance. The ability to accurately distinguish germline from somatic SVs in the absence of a matched normal will enable functional assessment of factors guiding SV formation and consequences for therapy development. Population-level databases such as gnomAD may contain somatic variants resulting from aging-related processes; this tool may provide for a more accurate catalog of variants.

## Methods and Materials

### Data Acquisition

Whole genome sequencing tumor samples with matched normals were obtained from the TCGA patient cohort (**Supplementary Table 3**)^27^. We used SvABA to call SVs on all samples as previously described, which were used for our training and test sets^10^. The pHGG test set consisted of six samples from our recent study^28^. We also ran SvABA on the pHGG samples, which called the SVs used in our external test set.

### Quality Control and Filtering

We obtained the filtered “sv.vcf” SvABA outputs for germline and somatic breakpoints from all samples of our cohort. We then grouped pairs of breakpoints according to their MATEID. Any breakpoint without an identified MATEID pair was removed. We will subsequently refer to these MATEID pairs as rearrangements (or SVs). Once combined, we selected SVs that had the max MAPQ value (60) for both breakpoints, were not detected solely by discordant reads, had a span of 1000bp or greater or were translocations, and had at least two SV-supporting split reads.

Once the TCGA rearrangements were filtered, two-thirds (555,849 SVs) were randomly selected to be part of the training set, and the remaining one-third (277,925 SVs) were labeled as the test set.

### gnomAD Filtering

We obtained the gnomAD v4.0 release from the open-source gnomAD browser^29^. We filtered for variants that passed all gnomAD filters and had resolved breakpoints. In addition, we excluded all complex (CPX) SVs due to unclear breakpoint origins, as well as insertion (INS) SVs that lacked a source sequence. For insertion SVs that reported a source sequence, we resolved the breakends to reflect a corresponding intrachromosomal or interchromosomal translocation.

### gnomAD Fuzzy Matching

For each rearrangement in our test set, we first determined the distance to the closest gnomAD reference SV. The distances of a given rearrangement breakpoint to its “corresponding” breakpoint of all reference gnomAD SVs were calculated on each side (i.e. comparing the 5’ breakpoint of the rearrangement to the 5’ breakpoint of a gnomAD reference SV and vice versa). These distances were averaged, and the gnomAD SV with the lowest average was labeled as the closest SV.

Each of these discrete values of average distances was treated as a cutoff, and we constructed an ROC curve by classifying all rearrangements with an average distance less than that cutoff as germline. For each of the cutoffs, the predicted and actual classifications were used to calculate the specificity and sensitivity, which were used to generate the ROC curve.

### Feature Annotation

Once the filtering step was complete, we annotated each SV with the following features: distance to the closest gnomAD reference SV, DNA replication timing of each breakpoint, GC content and length of any novel sequence insertion, GC content and length of homology associated with a breakpoint, TP53 gene mutation status, number of other SVs within a 5Mbp window, total number of SVs in the sample, distance to a long interspersed nuclear element (LINE), distance to a short interspersed nuclear element (SINE), distance to the nearest SV, span of the SV, type of the SV (categorized as deletion, duplication, inversion, or translocation), impact on a gene, and impact on an exon.

The distance to the closest gnomAD reference SV for a given rearrangement was encoded as two features, each representing the distance of a breakpoint to its “corresponding” breakpoint of the nearest reference gnomAD SV, similar to the method used for gnomAD fuzzy matching. If a matching gnomAD SV was not found, as is the case for many translocations, we input an artificial distance of 1e9 for each breakpoint.

The DNA replication timing was likewise encoded as two features, corresponding to the replication timing of each breakpoint. This was determined by looking up each breakpoint location in the DNA replication timing table for the hg19 reference genome (from http://mskilab.com/fishHook/hg19/RT_NHEK_Keratinocytes_Int92817591_hg19.rds).

The GC content and length of insertion and microhomology sequences were features derived from the SvABA output. When assembling the contigs, SvABA frequently finds short gaps or overlaps between regions that map to the reference genome. These gaps or overlaps were output as “INSERTION” or “HOMSEQ” respectively. The gaps (i.e. the short region of the sample genome that did not map to the reference genome) and overlaps were analyzed for GC content and length and each of these were added as features.

The TP53 mutation status was determined using the consensus SNV, MNV, and indel calls from TCGA samples available on the ICGC Data Portal^30,31^. These consensus calls were filtered for the TP53 gene and functional mutations (i.e. variants classified as “3’UTR”, “5’UTR”, “lincRNA”, “Intron”, “Silent”, “IGR”, “5’Flank”, “RNA” were removed). XX of our samples were not analyzed in the PCAWG consensus calls. For these, we assigned the TP53 status as indeterminate. Each sample was then assigned a value of 1, 0, or-1, which corresponded to TP53 mutant, indeterminate, or TP53 wild-type.

The number of SVs in a 5Mbp window and the total number of SVs in the sample were calculated by counting the number of rearrangements that were within 5Mbp of the SV and within the specific sample respectively. The distances to the nearest SINE and LINE events were calculated by taking the distance from a given rearrangement to the closest SINE and LINE events^32^. The distance to the nearest SV was calculated by iterating through other rearrangements in the same sample, determining the distance between the given rearrangement and the other SV, and selecting the lowest value.

The span of the SV was taken from the SvABA output. The SV-type feature was separated into four binary features, each corresponding to insertion, duplication, inversion, or translocation events based on the position and read orientation of the supporting reads

The impact on a gene or exon region were binary factors representing whether a given rearrangement overlapped a gene or exon region, which was downloaded from Gencode and Ensembl genome browser BioMart, respectively. If the SV breakpoint was located within one of these regions, it was labeled with a 1. If no overlap was found, the feature was labeled with a 0 (also see **Supp. Fig. S9A**).

Moreover, we log-transformed any feature that related to genomic distances or counts of SVs to reduce noise. These features included the SV span, homology length, insertion length, distance to the closest gnomAD reference SV, distance to the nearest SINE and LINE element, the number of SVs in a given sample, the distance to the nearest SV, and the number of SVs within 5Mbp.

Once all the features were added, each feature was then scaled. The scaling was done by creating a scaling matrix containing values representing the mean and standard deviation of each feature (**Supplementary Table 4**). These values were calculated by taking 10 random samples of 50,000 SVs from the training dataset and evaluating the mean and standard deviation of the features for all of these rearrangements. Each input rearrangement feature—including those in both training and test sets—was scaled by subtracting the mean and dividing by the standard deviation in this matrix.

### Statistical Analyses

We performed Kolmogorov-Smirnov tests to determine if the distributions of continuous features between germline and somatic SVs significantly differed. Our p-value significance threshold was set to 0.05. For features quantified in discrete variables, we used Chi-squared tests with a significance threshold of 0.05.

To assess the significance of associations between variables both within SV classes and across classes, we performed a variety of statistical tests. For continuous/continuous feature comparisons within germline and somatic SVs, we computed Spearman ⍴ and defined moderate to high correlation as |⍴| ≥ 0.5. We then obtained p-values for the Spearman ⍴ values using two-sided t-tests as described previously^33^. Next, we performed false discovery rate (FDR) multiple hypothesis correction with a q-value threshold of 0.05. To check whether these correlations significantly differed across germline and somatic SVs, we derived z-scores from the ⍴ values using Fisher’s z-transformation ^34^ and subsequently obtained p-values. We performed FDR correction on these p-values and defined significance at a q-value threshold of 0.05.

For continuous/discrete variables, we calculated Wilcoxon ranked sum tests. Within both somatic and germline SVs, we compared observations in the binary ingroup to those in the binary outgroup. We reported the Hodges Lehmann estimator of location shift and the FDR-corrected p-values with a significance threshold of 0.05. For across-group comparisons, we also computed Wilcoxon ranked sum tests. Here, we compared continuous variable observations in the binary ingroup of germline SVs against the binary ingroup of somatic SVs.

For binary/binary variables, we performed Chi-squared tests on 2×2 contingency tables within germline and somatic SVs. We only compared binary variables that were non-mutually exclusive to obtain a relevant statistic. We reported the odds ratio and FDR-corrected p-values with a significance threshold of 0.05. For across-group binary/binary comparisons, we implemented the Breslow-Day test for homogeneity of associations on the 2×2×2 contingency tables^35^. This test checks if the odds-ratio across different strata-in this case, germline and somatic-are significantly different for any pair of binary variables. We reported the difference in odds ratios across the different strata and FDR-corrected p-values with a 0.05 significance threshold.

### Logistic Regression

We trained 21 logistic regression models-one for each feature-using the glm method from the R stats package. Our training set included a random sample of 200k SVs from the full training set described in the Quality Control and Filtering section that were scaled as described in the Feature Annotation section. We used each single-feature logistic regression model to compute prediction probabilities on 100k scaled SVs sampled from the full test set. Prediction probabilities were derived using the R stats predict method. Finally, we calculated the AUC, TPR, and FPR values to evaluate the performance of each model.

### SVM

We used an SVM with an RBF kernel from the e1071 package in R and set cost and gamma parameters as 10 and 0.1 respectively. These hyperparameters were determined by a grid-search, of all combinations of the following values: cost of 1, 10, 100, 500, 1000 for both RBF and linear kernels and gamma of 0.0001, 0.001, 0.01, 0.05 0.1, 1 for the RBF kernel. For each combination of hyperparameters, we performed a 5-fold cross-validation, splitting the training set into five groups. One group was selected as a validation set for each of the five cross-validation iterations, and the other four groups made up the sub-training set. During each iteration, the model was trained on the sub-training set and evaluated on the validation set. The resulting AUC and PPV for all iterations were averaged for each combination of hyperparameters, and the model that performed well for both metrics was chosen.

Although SVMs typically do not have an inherent probability metric, there is a probability feature built into e1071 implementation of the SVM—based on Platt Scaling. We used this to generate the ROC curve for the AUC above and to select a probability cutoff that optimizes for our specific needs, instead of just accuracy. Because we aimed to reduce germline contamination for analysis of somatic variants, we decided on a cutoff that resulted in a high PPV.

To achieve this, we treated the probability cutoff as another hyperparameter and performed another 5-fold cross-validation for each probability value between 0 and 1 in 0.001 increments. This time, we optimized for the maximum TPR + PPV value, since we found that PPV itself was monotonically increasing with respect to the probability cutoff. We did not change the cost, gamma, and kernel type parameters in this probability tuning step, and were kept as 10, 0.1, and RBF kernel respectively.

In constructing our classifier, we chose a probability cutoff that optimized for the sum of TPR and PPV, to minimize germline contamination from rearrangements called somatic. However, the probability cutoff chosen for our classifier can be modified for other use cases, including those that demand correct germline calls or a high overall accuracy, in which case a cutoff of the maximum sum of the TNR and NPV or the maximum AUC in the same TCGA validation set can be used.

## Ancestry Analysis

Using consensus ancestry calls of TCGA patients from a previous study, we conducted a lookup of patients in our cohort to those in the consensus calls^36,37^. After excluding the 25 patients who were not in the consensus calls, the test set of 277,925 TCGA SVs—each with an associated patient—were labeled with their corresponding ancestry call. For our analysis, we did not consider any of the 50 admixed patients. We also did not include the “amr” and “sas”, corresponding to American and South Asian, ancestries, as there were only seven and two patients within those categories, respectively. For the remaining “afr”, eas”, and “eur”, corresponding to African, East Asian, and European, ancestries, we subsetted the test set according to their ancestry labels, and analyzed the performance of the classifier on each subset.

## Signature Analysis

The signature analysis was performed primarily using the <u>signature.tools.lib</u> package published by the Nik-Zainal group^3^. From bedpe files containing SVs for each patient in our cohort, we first created true germline and somatic signature catalogs for each patient using the `bedpeToRearrCatalogue` function in the signature.tools.lib package. These signature catalogs consisted of counts of the number of rearrangements organized across three categories: SV span, type, and clustering of all SVs of span above 1kb. This process was repeated for predicted germline and somatic rearrangements from both the fuzzy matching approach and the SVM classifier.

These catalogs were then fit to the reference signature profile for rearrangements from the same study using the `Fit` and `plotFit` functions^3,38^. This resulted in the true and predicted germline and somatic reference signature exposures for each patient, which shows how much contribution each reference signature has in the collection of SVs in each patient. For Fig 5A and Supplemental Fig S8, we determined the ratio of a given reference signature exposure to the sum of all exposures within a patient for all reference signatures. In the case that there were no SVs in a patient—for a given class—the proportion was simply set as 0, instead of an undefined value.

In Fig 5B, we first conducted a modified Spearman correlation between each predicted and true class. Each vector used for the Spearman correlation had a length of 974 and was composed of the proportion values for a given reference signature and a given class. Each predicted vector was paired with another true vector with the same reference signature. If both vectors were nonzero, then a normal Spearman correlation was conducted. If this was not the case, we calculated correlation as follows: If both vectors were zero,a Spearman rho of 1 was assigned. If one vector was zero and the other was nonzero, a Spearman rho of 0 was assigned.

## Availability of Data and Materials

All annotation functions, scripts used to create our figures, and trained classifier will be available on Github^39^. The datasets used to support the conclusions of this article are available online^27,29,31,32,37,38^. Any data generated during the current study are available from the corresponding author on reasonable request.

## Acknowledgments

We thank and acknowledge the following funding sources: Fund for Innovation in Cancer Informatics (S.D. and R.B.), the Gray Matters Brain Cancer Foundation (R.B.), the Pediatric Brain Tumor Foundation (R.B.), and Break Through Cancer (R.B.). F.D. is supported by the Max Eder program of the German Cancer Aid and a participant in the BIH Charité Junior Clinician Scientist program. S.D. is supported by an NIH NRSA award (F32), 1F32CA261024.

## Supplementary Figures and Figure Legends

**Supplementary Figure 1.**
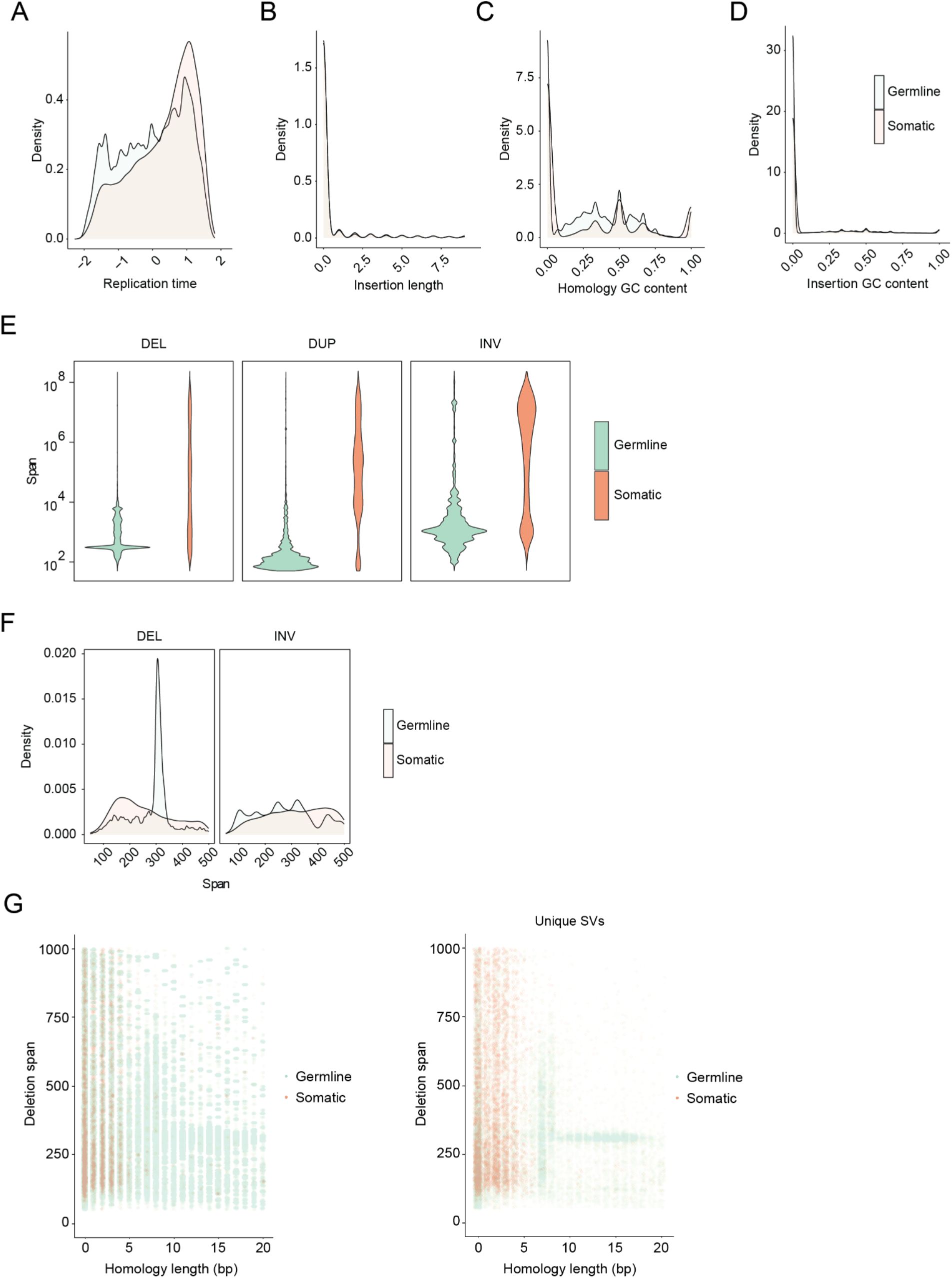
Germline and somatic distributions for additional features considered. **A,** Replication timing distributions for SV breakpoints. **B,** Length distribution of SV insertion sequence. **C,** GC content of homology sequence. **D,** GC content of insertion sequence. **E,** Violin plots of span distributions of germline and somatic deletions (DEL), duplications (DUP), and inversions (INV). **F,** Density plots of span distributions of deletions and inversions truncated at 500bp. **G,** Span of deletions vs homology lengths across the entire dataset (left) and downsampled to represent unique SVs (right) showing a peak in span distribution of germline deletions that coincides with homology lengths between 13-17 bp.

**Supplementary Figure 2.**
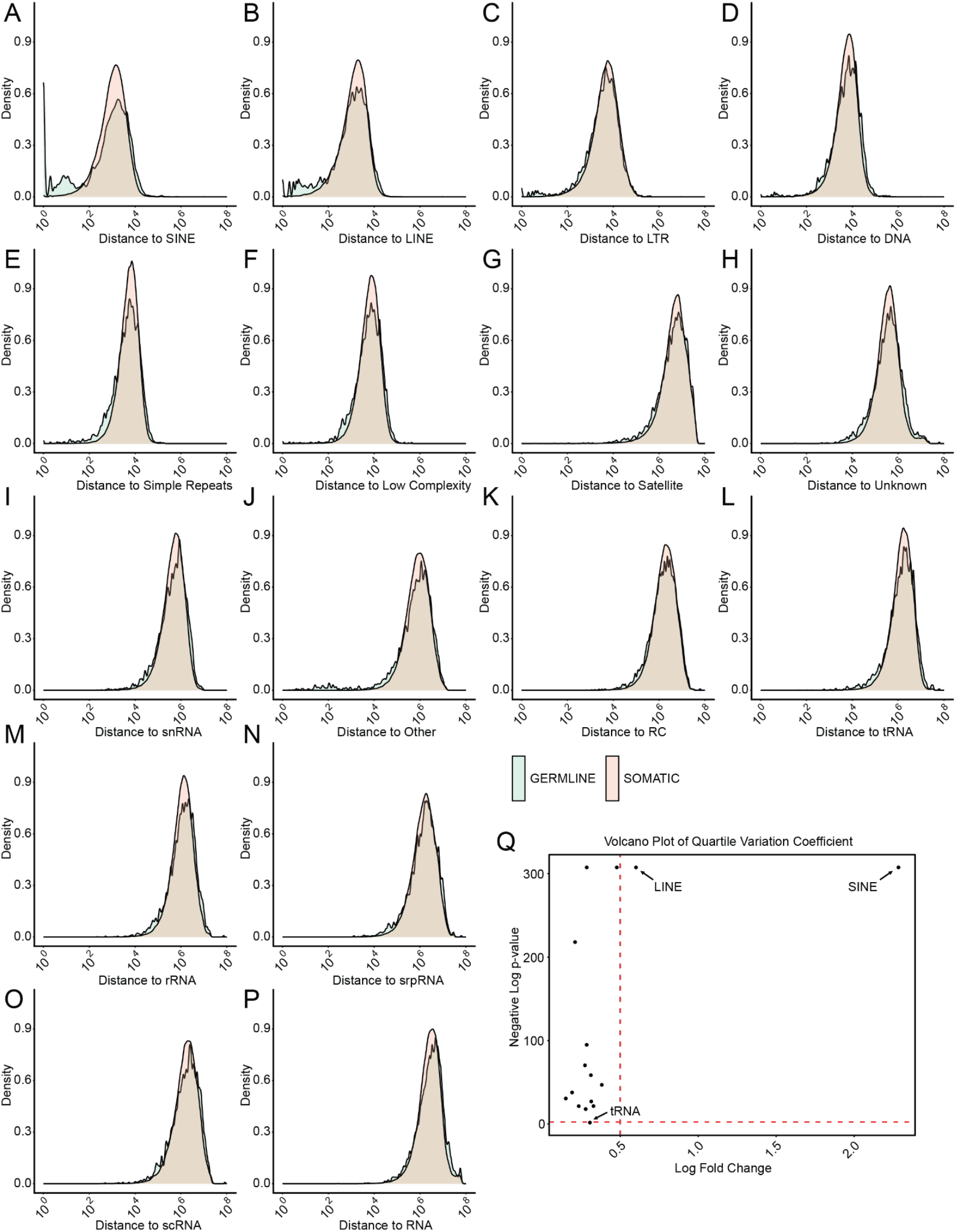
Distances to various repeat elements. **A-P,** Distributions of the distances of germline and somatic SVs to various repeat elements ordered by instances of each element in the genome. These include SINE (A), LINE (B), long terminal repeat (C), DNA (D), simple repeat (E), low complexity (F), satellite (G), unknown (H), snRNA (I), other (J), rolling circle (K). Distances to tRNA (L), rRNA (M), srpRNA (N), scRNA (O), RNA (P) pseudogenes were also calculated. **Q,** For each repeat element, a Wilcoxon test was performed, and the BH-corrected p-values were plotted in a volcano plot against the difference in quartile variation coefficient of the distributions. Only SINE and LINE elements passed both cutoffs.

**Supplementary Figure 3.**
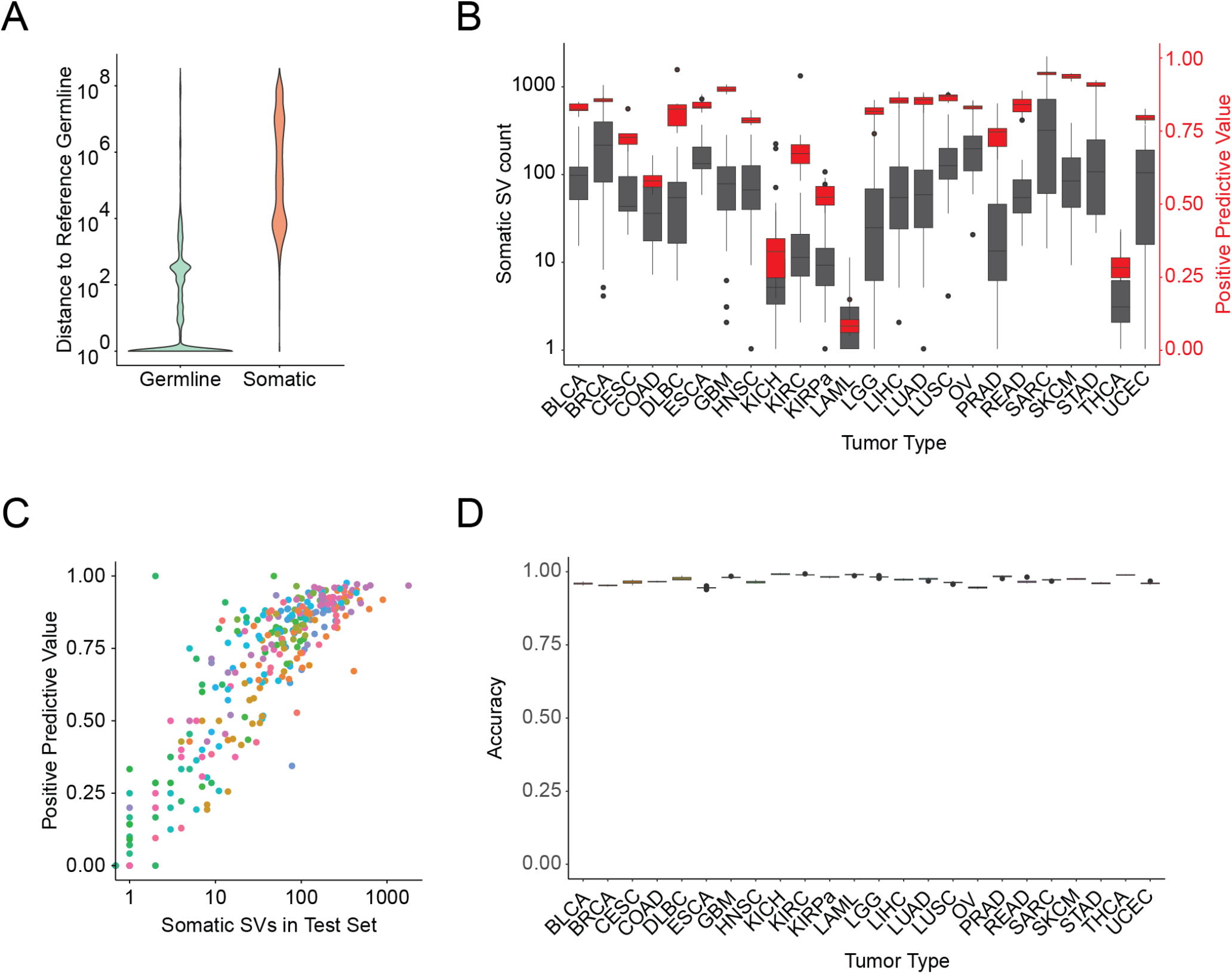
GaTSV classifier performance is related to somatic SV count. **A,** Distribution of total distance of germline and somatic SVs to the closest SV in gnomAD (reference) SV database. **B**, Somatic SV count, and PPV for each tumor type in the TCGA dataset. **C**, PPV vs. somatic SV count for every tumor in the TCGA dataset. **D,** Accuracy of GaTSV classifier across each tumor type

**Supplementary Figure 4.**
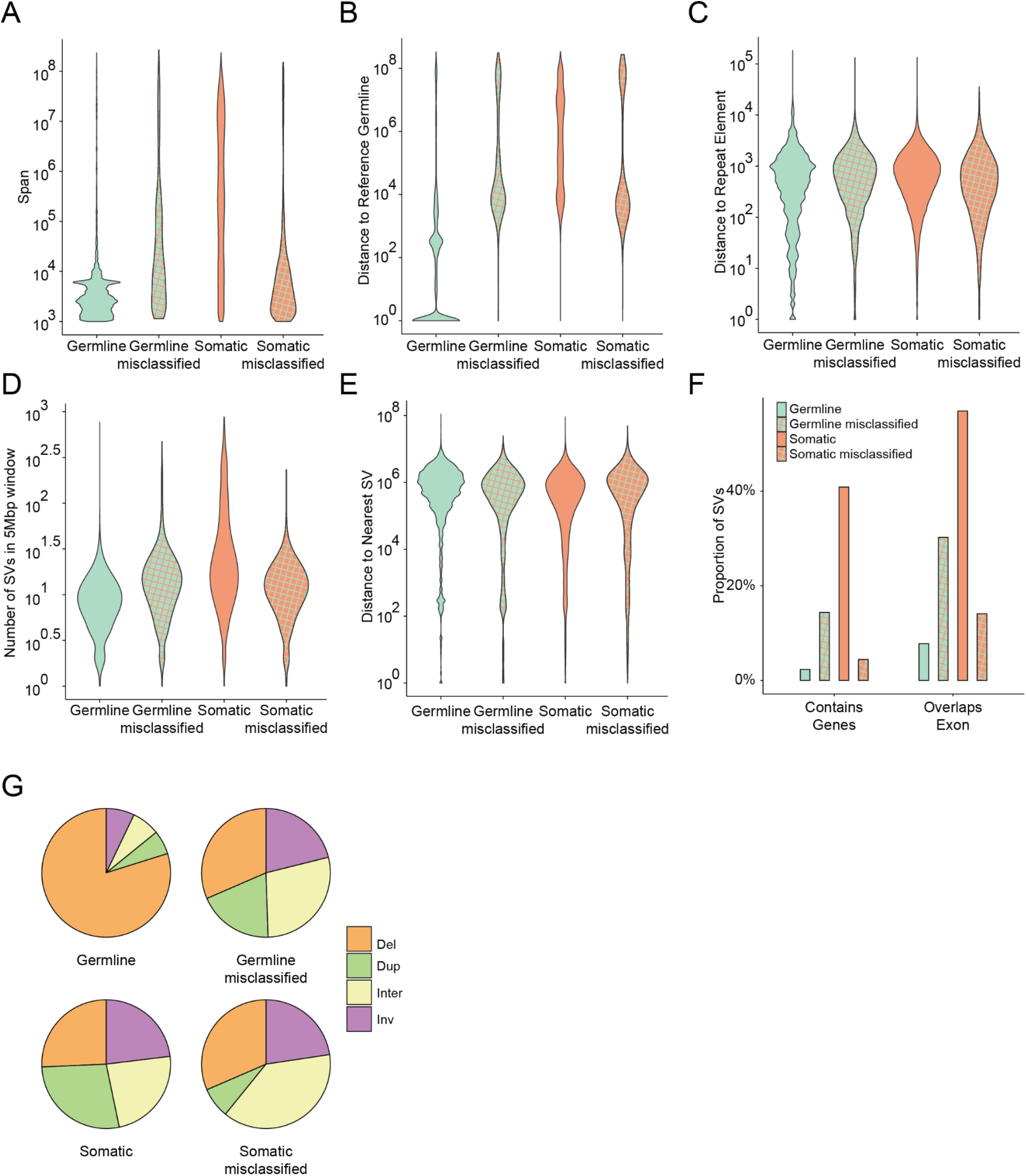
Misclassified SVs resemble the other class. Span distribution (**A**), distance to the closest SV in gnomAD (reference) database (**B**), distance to nearest repeat element (LINE or SINE) (**C**), number of SVs within a 5Mbp window (**D**), distance to nearest SV within a sample (**E**), proportion of SVs that impact a gene or overlap an exon (**F**), and proportion of each SV type-Del (deletion), Dup (duplication), Inter (interchromosomal/ translocation), Inv (inversion) (**G**) for correctly and incorrectly classified SVs.

**Supplementary Figure 5.**
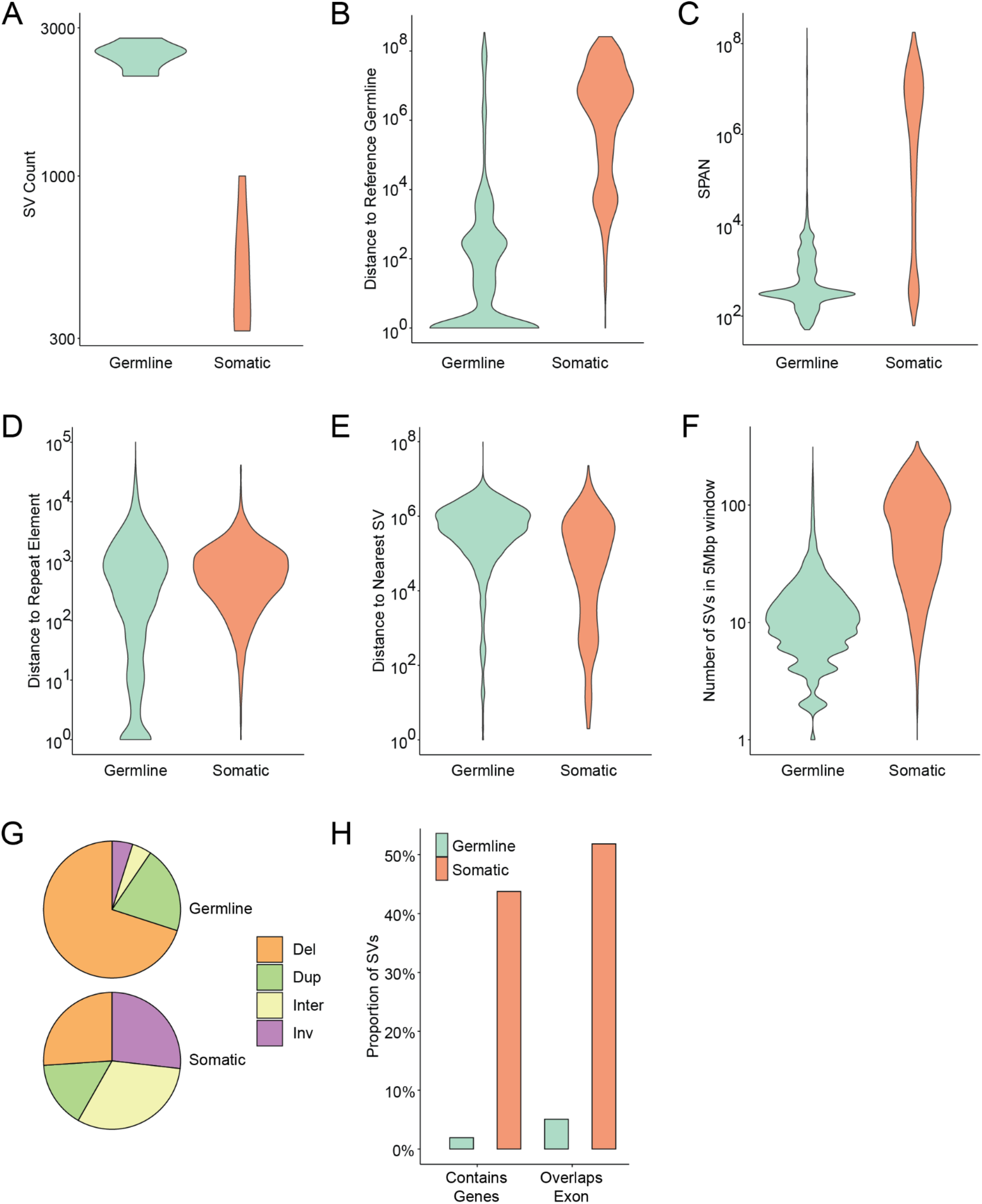
SV Feature Distribution in the pHGG test set is similar to TCGA. **A**, Germline, and somatic SV frequencies in test samples. **B**, Total distance of somatic and germline SVs to closest SV in gnomAD reference. **C**, Span distribution of somatic and germline SVs. **D**, Distance of somatic and germline SVs to nearest repeat element (LINE or SINE). **E**, Distance of germline and somatic SVs to the nearest SV within a sample. **F**, Number of SVs within a 5Mbp window of each SV breakpoint. **G**, Proportion of each SV type-Del (deletion), Dup (duplication), Inter (interchromosomal/ translocation), Inv (inversion) present in germline and somatic SVs. **H**, Proportion of germline and somatic SVs that impact a gene or overlap an exon.

**Supplementary Figure 6.**
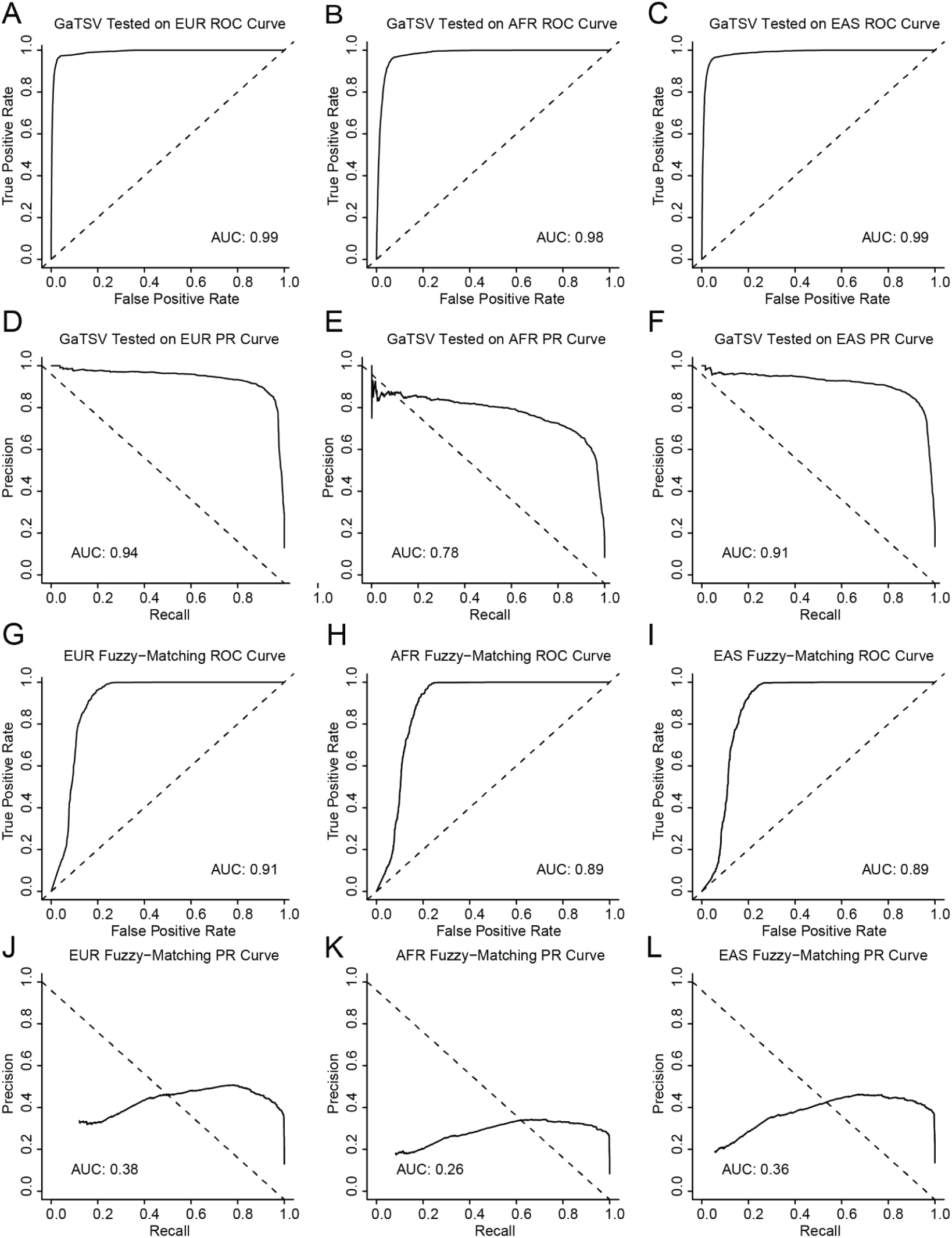
Classifier performances on ancestry-specific subsets of the TCGA test set. **A-C**, ROC for the GaTSV classifier on individuals descending from European (EUR) (A), African (AFR) (B), and East Asian (EAS) (C) ancestry. **D-F**, PR curve for the GaTSV classifier on individuals descending from European (D), African (E), and East Asian (F) ancestry. **G-I**, ROC for the fuzzy-matching method to the gnomAD database on individuals descending from European (G), African (H), and East Asian (I) ancestry. **J-L**, PR curve for the fuzzy-matching method on individuals descending from European(J), African(K), and East Asian(L) ancestry.

**Supplementary Figure 7.**
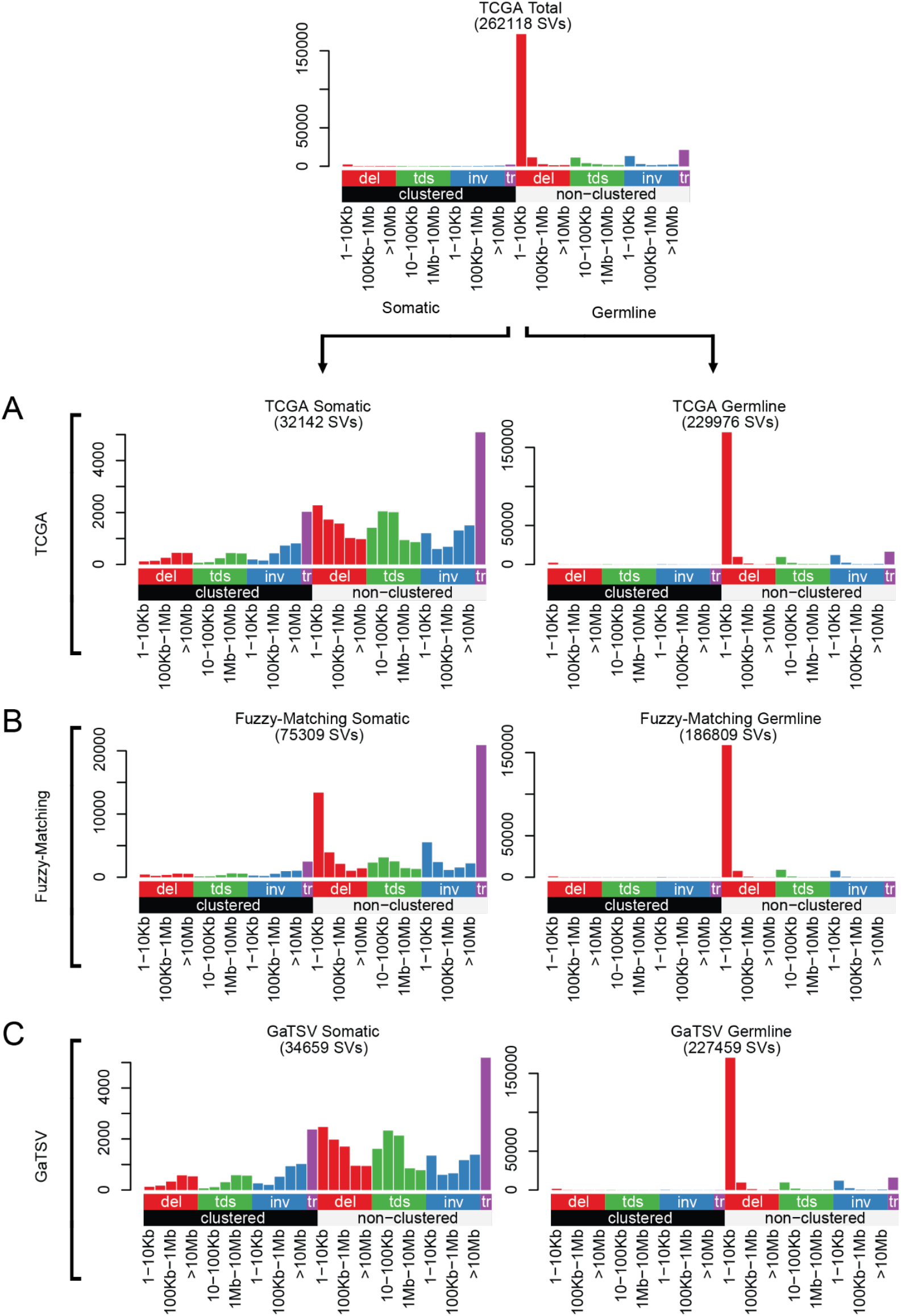
Distribution of SV categories (SV-Signature input catalogs) for true and predicted germline and somatic SVs. **A-C**, SV catalogs for the GaTSV classified SVs (C) match those of the true TCGA SVs (A) more closely than the fuzzy-matching to the gnomAD database (B).

**Supplementary Figure 8.**
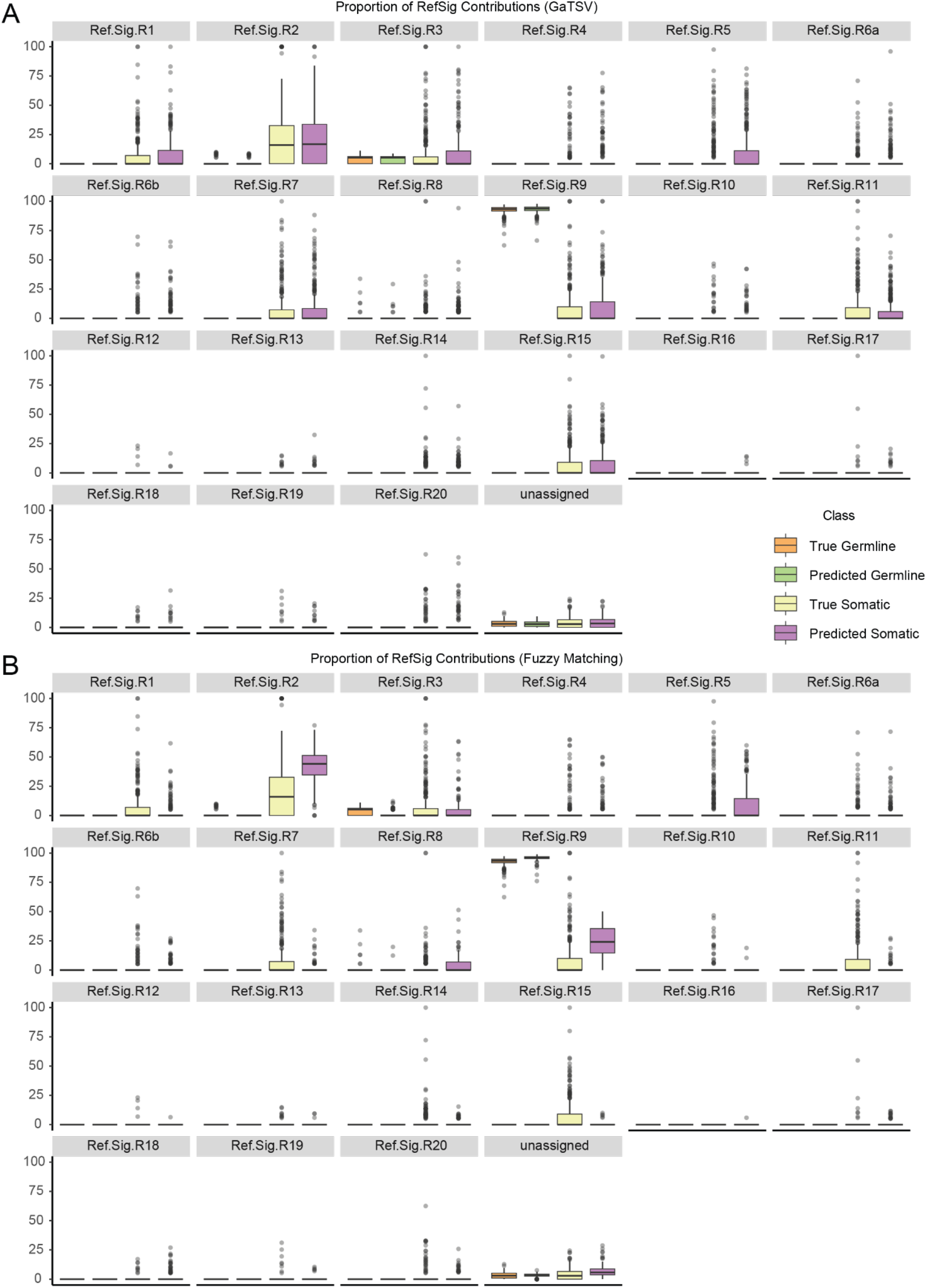
The proportion of individual SV signature contributions in each patient. **A**, post GaTSV classification. **B**, post fuzzy-matching to the gnomAD database.

**Supplementary Figure 9.**
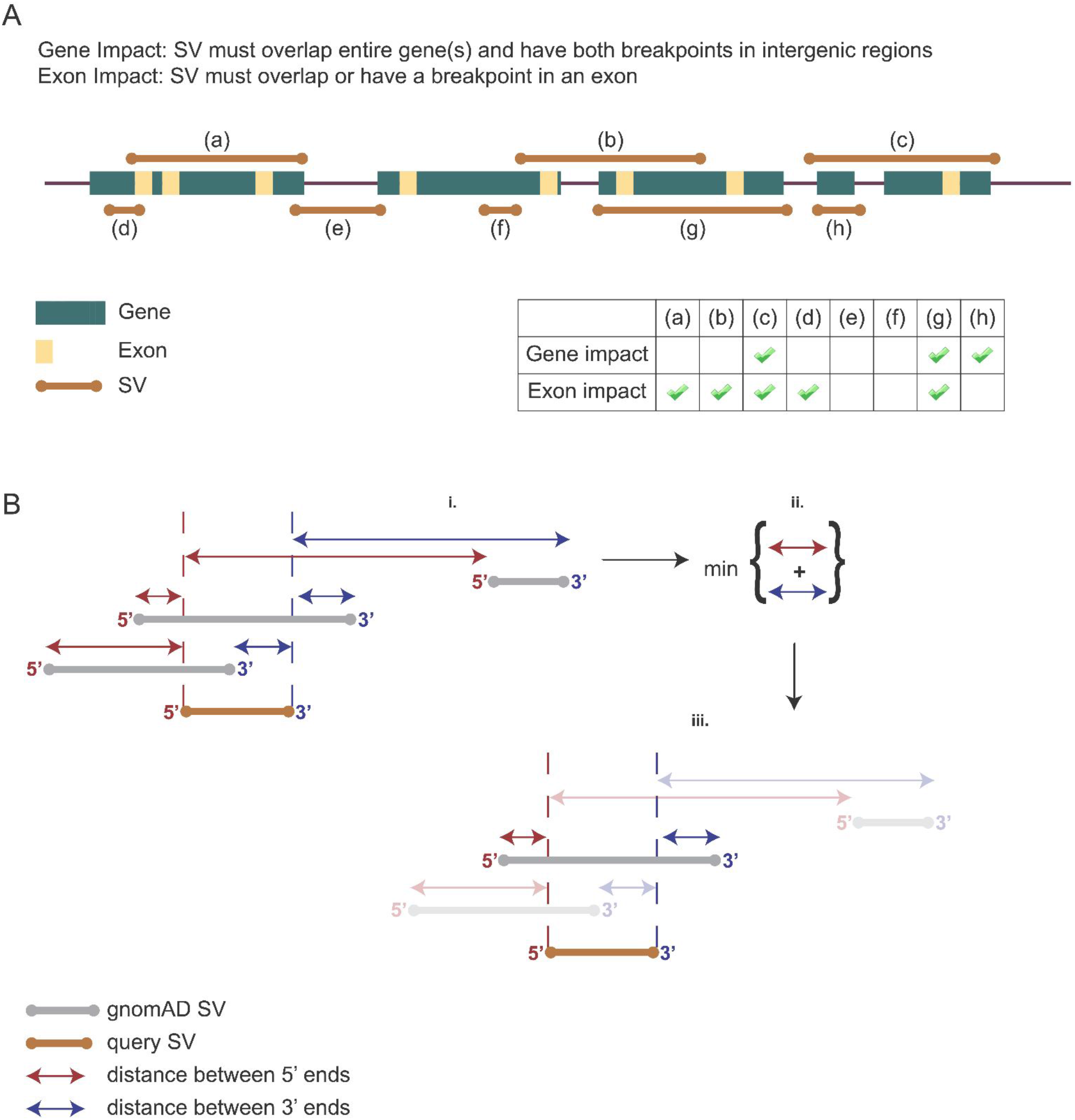
Schematics of how gene/exon impact was determined for non-translocation events (**A**) and how distance to nearest gnomAD SV was calculated (**B**).

## Supplementary Table Legends

**Supplementary Table 1.** Features of the SVM.

**Supplementary Table 2.** Performance of logistic regression models using single SV features.

**Supplementary Table 3.** TCGA cohort patient metadata.

**Supplementary Table 4.** Scaling matrix for SVM features.

