## Supplementary material for "Comparison of germline and somatic structural variants in cancers reveal systematic differences in variant generating and selection processes": Supp. Table 1

| Continuous Features | | Discrete Features | |
|---|---|---|---|
| **Code** | **Feature** | **Code** | **Feature** |
| homlen | Homology Length | del | Deletion |
| insertion_len | Insertion Length | dup | Duplication |
| SPAN | Span | inter | Interchromosomal event/translocation |
| gnomad_d_bkpt1 | Distance of left breakpoint to nearest SV in gnomAD | inv | Inversion |
| gnomad_d_bkpt2 | Distance of right breakpoint to nearest SV in gnomAD | CN_annot | Gene impact |
| hom_gc | % GC content in homology sequence | exon_annot | Exon impact |
| insertion_gc | % GC content in insertion sequence | tp53_status | TP53 mutation status (-1 (WT), 0 (Unknown), 1 (MUT)) |
| line_dist | Distance to nearest LINE | | |
| sine_dist | Distance to nearest SINE | | |
| sv_dist | Distance to nearest SV | | |
| sv_count_5Mbp | Total number of SVs in 5Mbp window of each SV breakpoint | | |
| sv_reptime_left | Replication timing of left breakpoint | | |
| sv_reptime_right | Replication timing of right breakpoint | | |
| num_sv_sample | Total number of SVs in the sample from which the SV under consideration is derived. | | |
| prop | % representation of the SV in TCGA pcawg cohort | | |

**Supplementary Table 1:** Features of the SVM
